# Transcutaneous auricular vagus nerve stimulation enhances emotional bias towards happiness in healthy young adults: A comparative study of electrical and ultrasound stimulation

**DOI:** 10.1101/2025.11.19.689293

**Authors:** June Christoph Kang, Ji You Ng, Marcus Kaiser, Hyuk Choi, Jae-Jun Song, JeYoung Jung

**Author notes:** Corresponding Author Dr JeYoung Jung, School of Psychology, University of Nottingham, University Park, Nottingham NG7 2RD UK, Dr June Christoph Kang, Department of Brain and Cognitive Engineering, Institute of Brain and Cognitive Engineering, 145 Anam-ro, Seoul, 02841, Republic of Korea.

## Abstract

Transcutaneous auricular vagus nerve stimulation (taVNS) is an emerging neuromodulation technique demonstrating promise in emotional regulation. This study investigated the acute effects of both electrical (E-taVNS) and ultrasound (U-taVNS) modalities on emotional bias using a facial emotion categorization task in healthy young adults. Fifty-nine participants underwent a single-blind, sham-controlled, within-subject design, with emotional bias assessed at pre-, during-, and post-stimulation phases. Our findings revealed that both E-taVNS and U-taVNS significantly enhanced emotional bias scores, shifting the perception of neutral and ambiguous faces towards positive interpretations and reducing negative emotional bias. No significant differences in efficacy were observed between the two stimulation modalities.

Furthermore, individual differences in interoceptive awareness were found to be associated with the observed taVNS effects. These results suggest that both electrical and ultrasound taVNS can acutely modulate emotional regulation, highlighting the potential of taVNS as a non-invasive, well-tolerated alternative for interventions targeting emotional bias and mood disorders.

**Highlights:**

- taVNS reduces negative emotional bias toward ambiguous stimuli.
- Both electrical and ultrasound taVNS shifted emotional bias toward more positive interpretations of ambiguous facial expressions.
- Ultrasound taVNS (U-taVNS) demonstrated comparable efficacy to electrical taVNS (E-taVNS).
- The subscale of the Multidimensional Assessment of Interoceptive Awareness predicted individual variability in taVNS-induced emotional bias change.

## Introduction

Emotion recognition is fundamental to social interaction (Calvo & Nummenmaa, 2016), enabling rapid evaluation of social environments, appropriate attention allocation, and interpretation of ambiguous social cues (Bargh et al., 2012). Disruptions to this adaptive process are common in mood and anxiety disorders (Surguladze et al., 2004; Joormann & Gotlib, 2006). In particular, negative emotional biases such as heightened attention to threat in anxiety or a tendency to interpret ambiguous stimuli negatively or recall negative over positive experiences are well documented (Hayes & Hirsch, 2007; Joormann & Gotlib, 2006). Accordingly, interventions that shift maladaptive biases toward more adaptive interpretations hold promise for both clinical and healthy populations. Behavioural and non-invasive brain stimulation approaches, including transcranial magnetic stimulation (TMS) and transcranial electrical stimulation (tES) have targeted emotional bias to reduce depression and anxiety symptoms (Briley et al., 2024; Coussement et al., 2019; Nord et al., 2017; Suddell et al., 2023). While some studies reported positive outcomes (Briley et al., 2024; Suddell et al., 2023), others yielded inconsistent findings (Coussement et al., 2019; Nord et al., 2017). In contrast, transcutaneous vagus nerve stimulation (taVNS) modulates emotional processes indirectly through vagal pathways rather than direct cortical stimulation, and has shown promising therapeutic effects in anxiety major depressive disorder (Austelle et al., 2025)

The vagus nerve, a central component of the parasympathetic nervous system, regulates autonomic balance, affective and interoceptive processes. Roughly 80% of its fibers are afferent, transmitting sensory information to the nucleus tractus solitarius (NTS) in the brainstem, which projects to emotion-related regions including the amygdala, hypothalamus, insula, and nucleus accumbens (Bonaz et al., 2018; Wang et al., 2021; Butt et al., 2020; Yakunina et al., 2017).

Traditionally, invasive VNS involved surgically implanted electrodes around the cervical vagus nerve (Groves & Brown, 2005), but its application is limited by complications such as hoarseness and infection (Yap et al., 2020). taVNS offers a non-invasive alternative, delivering electrical stimulation to the auricular branch of the vagus nerve (ABVN), typically via electrodes placed on the tragus or cymba conchae of the external ear (Hilz, 2022).

Neuroimaging studies showed that taVNS activated the same central circuits as invasive VNS, including the NTS, locus coeruleus (LC), amygdala, and prefrontal cortex (Frangos et al., 2015; Kraus et al., 2007; Yakunina et al., 2017). A key mechanism involves modulation of the LC-norepinephrine (LC-NE) system, which regulates arousal, attention, and adaptive behaviour (Colzato & Beste, 2020; Ghosh & Maunsell, 2024). By increasing phasic noradrenergic activity, taVNS may enhance cognitive control and emotional regulation. Physiological markers such as pupil dilation (Ludwig et al., 2024; Skora et al., 2024), and P300 event-related potentials (Ventura-Bort et al., 2021), and salivary alpha-amylase (Giraudier et al., 2022) further support LC-NE engagement. The enhanced LC-prefrontal connectivity has also been associated with more positive interpretation of ambiguous facial expressions (Dave et al., 2025).

taVNS also influences amygdala–prefrontal connectivity, a pathway critical for balancing bottom-up emotional reactivity with top-down regulation (Liu et al., 2016). In addition, taVNS affects broader autonomic functions including cardiacrespiratory, and immune regulation (Badran et al., 2022; Bonaz et al., 2016). Empirical evidence showed that taVNS reduced depressed mood and negative emotional states (Zhao et al., 2025), and enhanced mood recovery after prolonged stress exposure (Ferstl et al., 2022). Its association with parasympathetic activity also supports positive mood regulation (Koenig et al., 2021; Yuan & Silberstein, 2016) and improved emotional inhibitory control via prefrontal networks, including the orbitofrontal and dorsolateral prefrontal cortex (Zhu et al., 2024). Consistent with evidence that taVNS shifts emotional bias in a positive direction, Osnes et al (2023) demonstrated that individuals with lower heart rate variability (HRV), a marker of vagal tone exhibited a stronger negativity bias, while those with higher HRV tended to interpret negative stimuli more positively Recent advances in neuromodulation have introduced low-intensity focused ultrasound (LIFU) as a promising non-invasive method for stimulating the auricular branch of the vagus nerve (Hacker et al., 2023; Kohler et al., 2025). Unlike traditional electrical transcutaneous auricular vagus nerve stimulation (E-taVNS), ultrasound neuromodulation operates through mechanical energy to induce pressure changes between intracellular and extracellular fluid, resulting in membrane curvature changes and action potential propagation without the discomfort often associated with electrical stimulation (Dell’Italia et al., 2022; Riis & Kubanek, 2021; Yoo et al., 2022). Preliminary evidence demonstrates that ultrasound taVNS (U-taVNS) can reduce symptoms of anxiety, depression, and PTSD, showing substantial improvements within weeks (Kohler et al., 2025). Given the vagus nerve’s projections to the amygdala and hippocampus, key structures for emotional processing and extinction learning (Noble et al., 2017), U-taVNS may offer comparable modulation of emotional processing with enhanced comfort and compliance (Kaniusas et al., 2019)

In this study, we investigated the effects of E-taVNS and U-taVNS on emotional bias in healthy adults. Participants completed a facial emotion categorization task with ambiguous expressions systematically morphed along a sad-happy continuum. Before the experiment, participants completed questionnaires assessing anxiety, depression, and interoceptive perception. Each participant received either E-taVNS or U-taVNS under active and sham conditions, performing the emotion categorization task before, during, and after stimulation. We hypothesized that both active E-taVNS and U-taVNS would enhance positive emotional bias relative to pre-stimulation and sham conditions. Furthermore, we compared the efficacy of E-taVNS and U-taVNS and explored whether baseline traits were associated with individual variability in taVNS effects.

## Method

### Participants

An a-priori sample size calculation was performed using G*Power 3.1.9.7 (Faul et al., 2009) to estimate the approximate number of participants required for a within-subject design (power= 0.8, alpha= 0.05). Based on a moderate effect size based on previous study (Johnson & Steenbergen, 2022), the estimated sample size was 19 participants. To ensure sufficient power and account for potential drop-out, 30 participants were targeted per stimulation modality.

In total, 59 healthy adult volunteers (46 females and 13 males, M= 23.6, SD= 2.88) were recruited. Participants were randomly assigned to either electrical stimulation (E-taVNS; n=30) and ultrasound stimulation (U-taVNS; n=29).

All participants completed the VNS Safety Questionnaire prior to enrollment to ensure eligibility. Exclusion criteria included: age under 18; any active implantable medical device (e.g., pacemaker, cochlear implant); history of carotid atherosclerosis or cervical vagotomy; cardiovascular conditions (hypertension, hypotension, bradycardia, tachycardia); metallic implants near the head or neck; current or past neurological or psychiatric disorders; predisposition to fainting; family history of epilepsy or seizures; current use of psychoactive medication (except hormonal contraceptives); or pregnancy. These criteria ensured participant safety and data integrity during non-invasive vagus nerve stimulation.

All participants provided written informed consent to participate in the study. The study was approved by the ethics committee of the University of Nottingham (F1619R) and conducted in accordance with the Declaration of Helsinki.

### Experimental design and procedures

Prior to the experiment, participants completed questionnaires, including the Beck Anxiety Inventory (BAI; Beck et al., 1988), the Beck Depression Inventory (BDI; Beck et al., 1996), and the Multidimensional Assessment of Interoceptive Awareness (MAIA-2; Mehling et al., (2018)).

Each experimental session (active and sham) followed a structured timeline comprising three phases: pre-stimulation, during stimulation, and post-stimulation (Figure 1a). Sessions were separated by a minimum of five days, and the order of stimulation was counterbalanced across participants.

**Figure 1.**
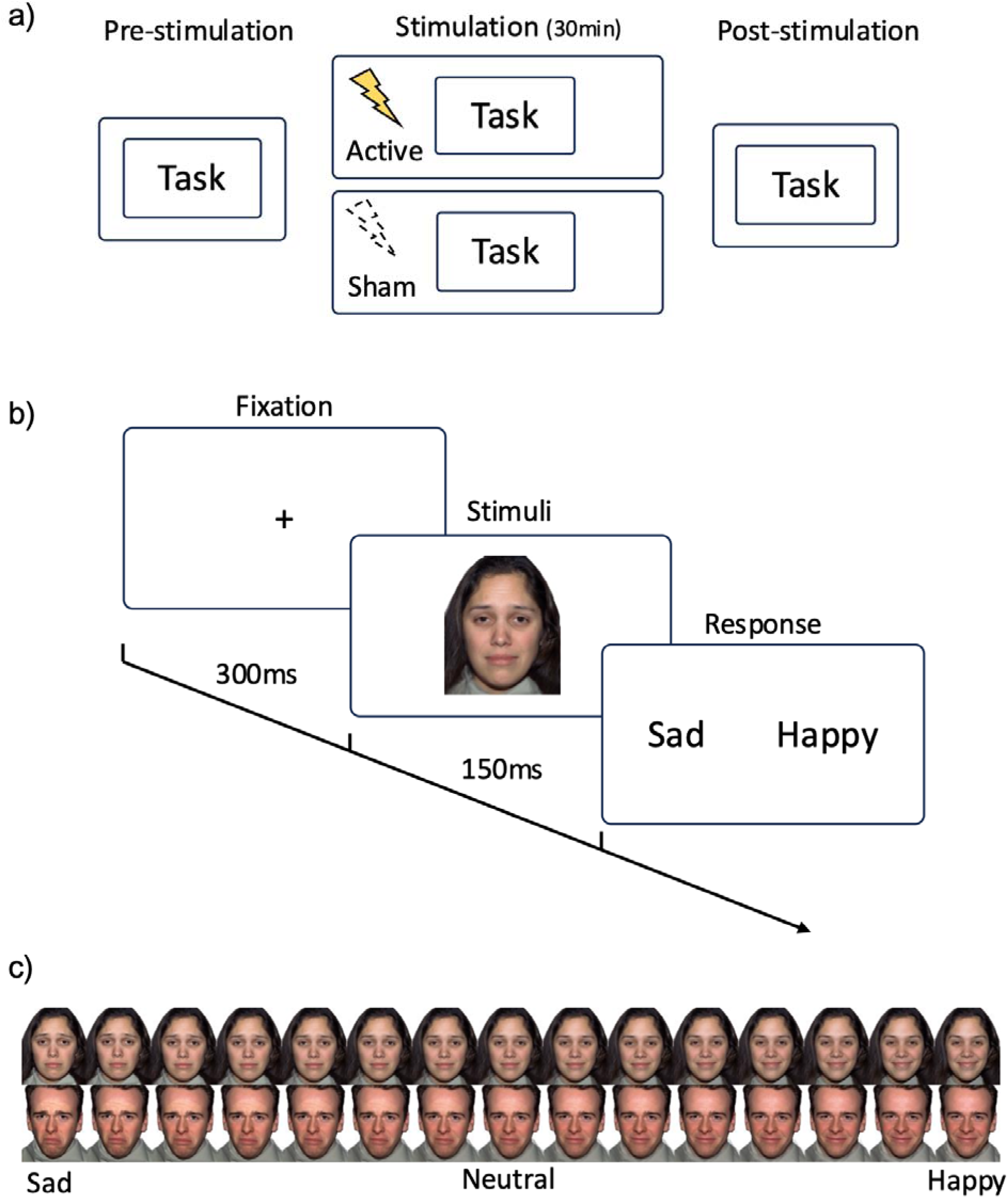
Overview of Experimental Paradigm and Stimuli. (a) Schematic overview of the experimental structure, illustrating the sequence of pre-stimulation, during-stimulation, and post-stimulation assessments. (b) Timeline of the emotional bias task showing trial structure, including fixation, brief presentation of a morphed facial stimulus, and subsequent response screen. (c) Examples of morphed facial stimuli used in the task, depicting a graded continuum from sad to happy expressions.

In each session, participants first completed the emotional bias task, categorizing morphed facial expressions along a sad–happy continuum (Figure 1b). Immediately afterward, taVNS was administered for 30 minutes. The initial 15 minutes served as a stabilization period to allow stimulation effects to emerge, after which participants repeated the emotional bias task during stimulation. Following the 30minute stimulation period, participants performed the task a final time (post-stimulation phase) (Figure. 1a). At the end of each session, participants completed a side-effects questionnaire to report any mild adverse sensations (e.g., headache, neck discomfort, nausea, tingling, or muscle contractions). After completing both sessions, all participants were debriefed and reimbursed for their time.

### Emotional Bias Task

The emotional bias task was programmed and administered using PsychoPy (v2024.2.1) to assess participants’ perceptual sensitivity and bias towards sad and happy facial expressions. Each trial began with a 500ms fixation cross presented centrally on a grey background, followed by a 150ms presentation of a single morphed facial stimulus (Figure 1b). Immediately after, participants viewed a two-alternative forced choice screen and indicated via key press whether the face appeared “sad” or “happy”. Participants were instructed to respond as quickly and accurately as possible. Each session consisted of 120 trials. Trials were randomized to prevent any systematic bias from stimulus order. The primary outcome was the emotional bias score, calculated as the proportion of faces categorized as “happy” across trials. Higher scores reflected a positivity bias, whereas lower scores reflected a negativity bias.

Facial stimuli were selected from the NimStim Set of Facial Expressions (Tottenham et al., 2009). The experimental stimuli comprised morphed facial expressions representing a parametric continuum between prototypical happy and sad emotional expressions (Figure 1c). For each facial identity (F03, F07, M24, M34), a 15-level morph sequence was generated through linear interpolation of facial features, creating gradual transitions from 100% sad (morph level 1) through emotionally ambiguous expressions (morph level 8) to 100% happy (morph level 15).

This morphing procedure produced a systematic gradient of emotional intensity, with intermediate expressions exhibiting progressively greater perceptual ambiguity. To control for potential confounding variables, all stimuli underwent standardization procedures. Images were converted to grayscale to eliminate color-based processing differences and maintain consistent identity information across the emotion continuum. The stimulus set included four facial identities (two male, two female) from the original database.

### Transcutaneous auricular Vagus Nerve Stimulation (taVNS)

The transcutaneous auricular vagus nerve stimulation was administered using two modalities: electric current (using Healaon pro, Neurive Inc., Gimhae, Republic of Korea) and ultrasound (using ZenBud, NeurGear Inc., Rochester, NY, USA) (Figure. 2).

**Figure 2.**
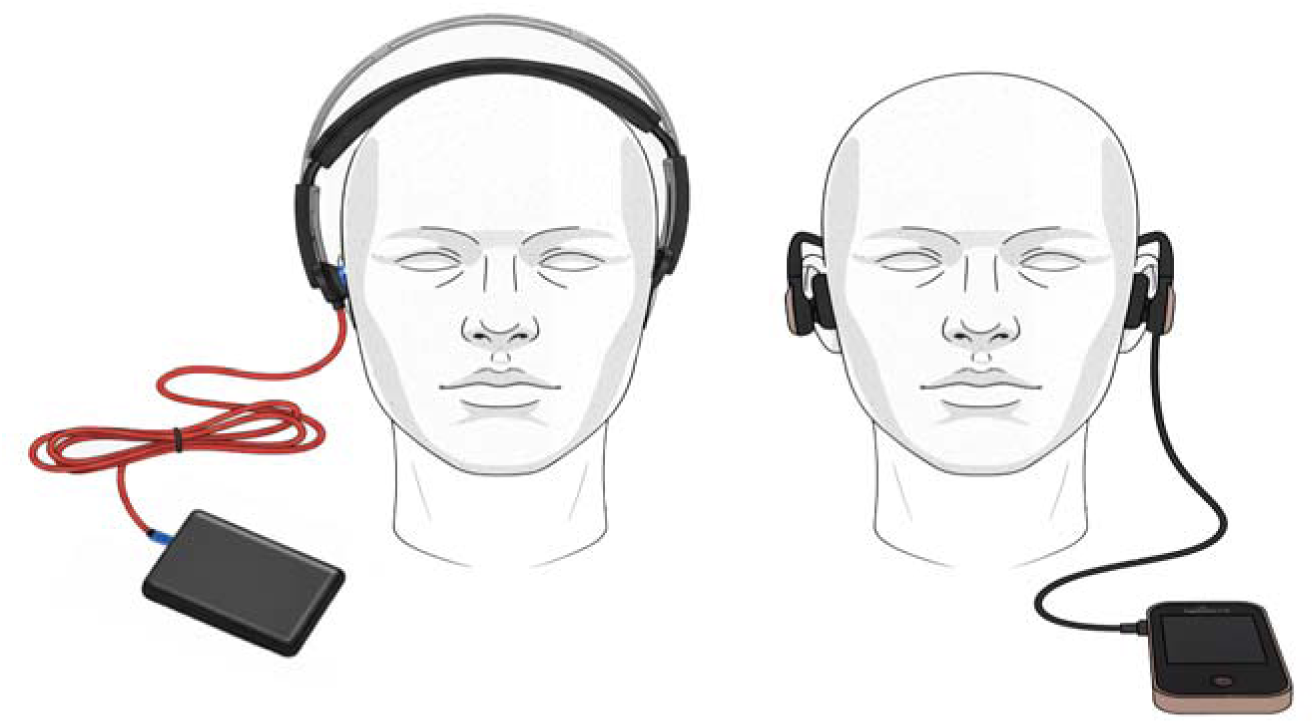
Stimulation Devices Used in the Study. Photographs of the ultrasound taVNS (U-taVNS) device (left) and the electrical taVNS (E-taVNS) device (right). The U-taVNS device delivers low-intensity focused ultrasound to the auricular branch of the vagus nerve, whereas the E-taVNS device delivers bilateral electrical stimulation to the cymba conchae.

For E-taVNS, stimulation was applied bilaterally to cymba conchae using conductive rubber electrode. The stimulation parameters were set at a frequency of 30 Hz and intensity level 5 (2.5 mA), with each session lasting 30mins. A continuous, wind-like auditory background was played throughout both active and sham conditions to mask any subtle device noise.

For U-taVNS, stimulation was delivered to the right cymba conchae. The ultrasound parameters included a centre frequency of 5.3 MHz, a pulse repetition frequency of 41 Hz, a duty cycle of 50%, a peak mechanical output of 0.081 W/ cm^2^,center frequency of 5.3 MHz, and an average intensity of 1.03 MPa.

### Statistical Analyses

All statistical analyses were performed in R (version 4.4.2). To examine the effects of Group (E-taVNS vs. U-taVNS), Stimulation (Active vs. Sham), and Phase (Pre, During, Post) on emotional bias scores, a three-way mixed-design ANOVA was performed. Post hoc pairwise comparisons assessed changes between pre-, stim, and post-stimulation phases.

To evaluate taVNS effects directly, delta bias scores (Post − Pre) were computed. Planned one-sample t-tests assessed whether delta scores differed from zero, and planned paired t-tests (one-tailed) compared delta scores between active and sham conditions. Independent t-tests compared the efficacy between E-taVNS and U-taVNS groups.

To examine adverse effects, a series of chi-square tests were conducted to compare frequencies of reported side effects across stimulation conditions (active vs. sham). All results are considered significant if p <.05.

## Results

### Demographics results

To ensure that the two groups were comparable prior to the experimental manipulation, independent-sample t-tests and chi-square tests were conducted on demographic and psychological variables (Table 1.). There were no significant differences between the e-taVNS and U-taVNS groups in sex distribution (X^2^(1, N = 59) = 0.00, p = 1.000), or in age (t(57) = - 0.28, p =.781). Similarly, no group differences were found in anxiety (BAI: t(57) =-0.47, p =.637; STAI-T: t(57) = 0.52, p =.603), depression (BDI: t(57) = 0.08, p =.939), or autism traits (t(57) =-0.04, p =.969). The two groups did not differ significantly in interoceptive awareness as measured by the MAIA (t(57) =-0.74, p =.463). No significant differences were found across any of the MAIA subscores, including Noticing (t(57) =-1.06, p =.292), Not-Distracting (t(57) = 0.17, p =.865), Not-Worrying (t(57) =-0.06, p =.953), Attention Regulation (t(57) =-1.26, p =.214), Emotional Awareness (t(57) =-0.44, p =.659), Self-Regulation (t(57) =-0.57, p =.574), Body Listening (t(57) = 0.26, p =.799), and Trusting (t(57) =-0.68, p =.501).

**Table 1.**
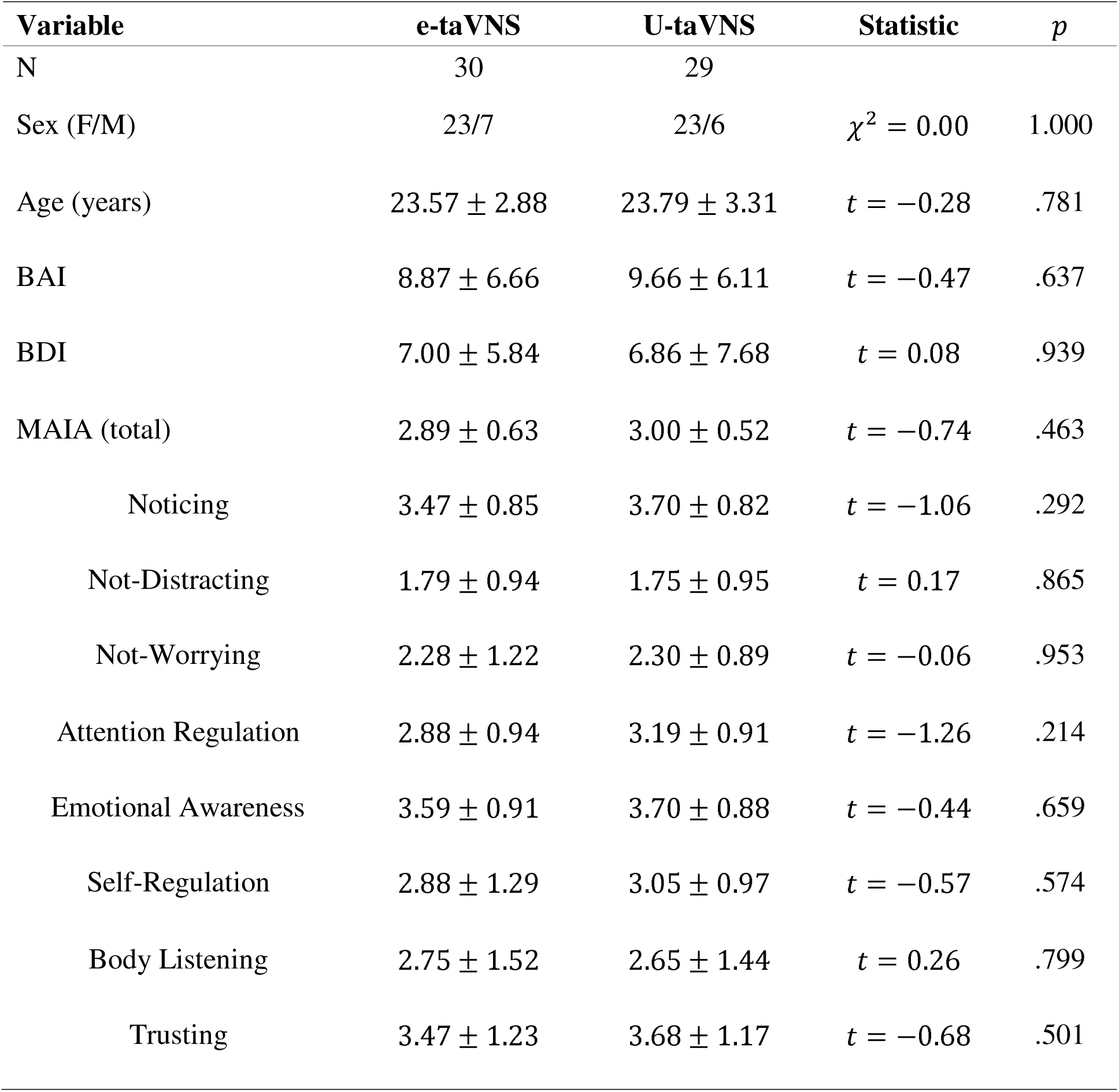
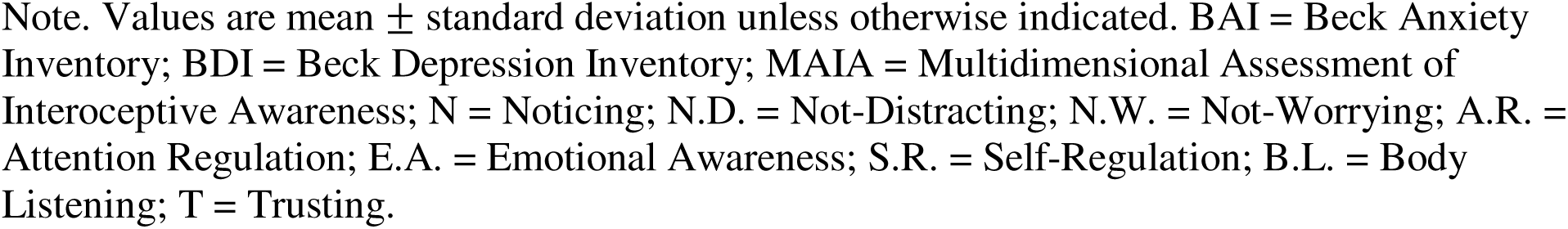
Participant Demographic and Psychological Characteristics. Group-level demographic and baseline psychological measures for participants assigned to the electrical taVNS (E-taVNS) and ultrasound taVNS (U-taVNS) conditions.

| Variable | e-taVNS | U-taVNS | Statistic | *p* |
| --- | --- | --- | --- | --- |
| N | 30 | 29 |  |  |
| Sex (F/M) | 23/7 | 23/6 | $\chi^2 = 0.00$ | 1.000 |
| Age (years) | $23.57 \pm 2.88$ | $23.79 \pm 3.31$ | $t = -0.28$ | .781 |
| BAI | $8.87 \pm 6.66$ | $9.66 \pm 6.11$ | $t = -0.47$ | .637 |
| BDI | $7.00 \pm 5.84$ | $6.86 \pm 7.68$ | $t = 0.08$ | .939 |
| MAIA (total) | $2.89 \pm 0.63$ | $3.00 \pm 0.52$ | $t = -0.74$ | .463 |
| Noticing | $3.47 \pm 0.85$ | $3.70 \pm 0.82$ | $t = -1.06$ | .292 |
| Not-Distracting | $1.79 \pm 0.94$ | $1.75 \pm 0.95$ | $t = 0.17$ | .865 |
| Not-Worrying | $2.28 \pm 1.22$ | $2.30 \pm 0.89$ | $t = -0.06$ | .953 |
| Attention Regulation | $2.88 \pm 0.94$ | $3.19 \pm 0.91$ | $t = -1.26$ | .214 |
| Emotional Awareness | $3.59 \pm 0.91$ | $3.70 \pm 0.88$ | $t = -0.44$ | .659 |
| Self-Regulation | $2.88 \pm 1.29$ | $3.05 \pm 0.97$ | $t = -0.57$ | .574 |
| Body Listening | $2.75 \pm 1.52$ | $2.65 \pm 1.44$ | $t = 0.26$ | .799 |
| Trusting | $3.47 \pm 1.23$ | $3.68 \pm 1.17$ | $t = -0.68$ | .501 |

Note. Values are mean ± standard deviation unless otherwise indicated. BAI = Beck Anxiety Inventory; BDI = Beck Depression Inventory; MAIA = Multidimensional Assessment of Interoceptive Awareness; N = Noticing; N.D. = Not-Distracting; N.W. = Not-Worrying; A.R. = Attention Regulation; E.A. = Emotional Awareness; S.R. = Self-Regulation; B.L. = Body Listening; T = Trusting.

### Effect of taVNS on Emotional Bias

A significant main effect of Condition was found, F(1,57) = 5.40, p =.023, 1]^2^ =.008 (Figure 3, Table S1), indicating greater reductions in negative emotional bias during active versus sham stimulation. There was also a significant main effect of Phase, F(2,114) = 14.43, p <.001, 1]^2^ =.023, reflecting changes across pre-, during-, and post-stimulation assessments. However, there were no significant effect of Group (p =.716), nor the interaction between Group, Condition, and Phase (all ps >.05). Post hoc comparisons revealed significant increases of bias score from pre to post stimulation in both the E-taVNS active condition (mean difference =-0.036, p= 0.032) (Figure 3a) and U-taVNS active condition (mean difference =-0.042, p = 0.005) (Figure 3b).

**Figure 3.**
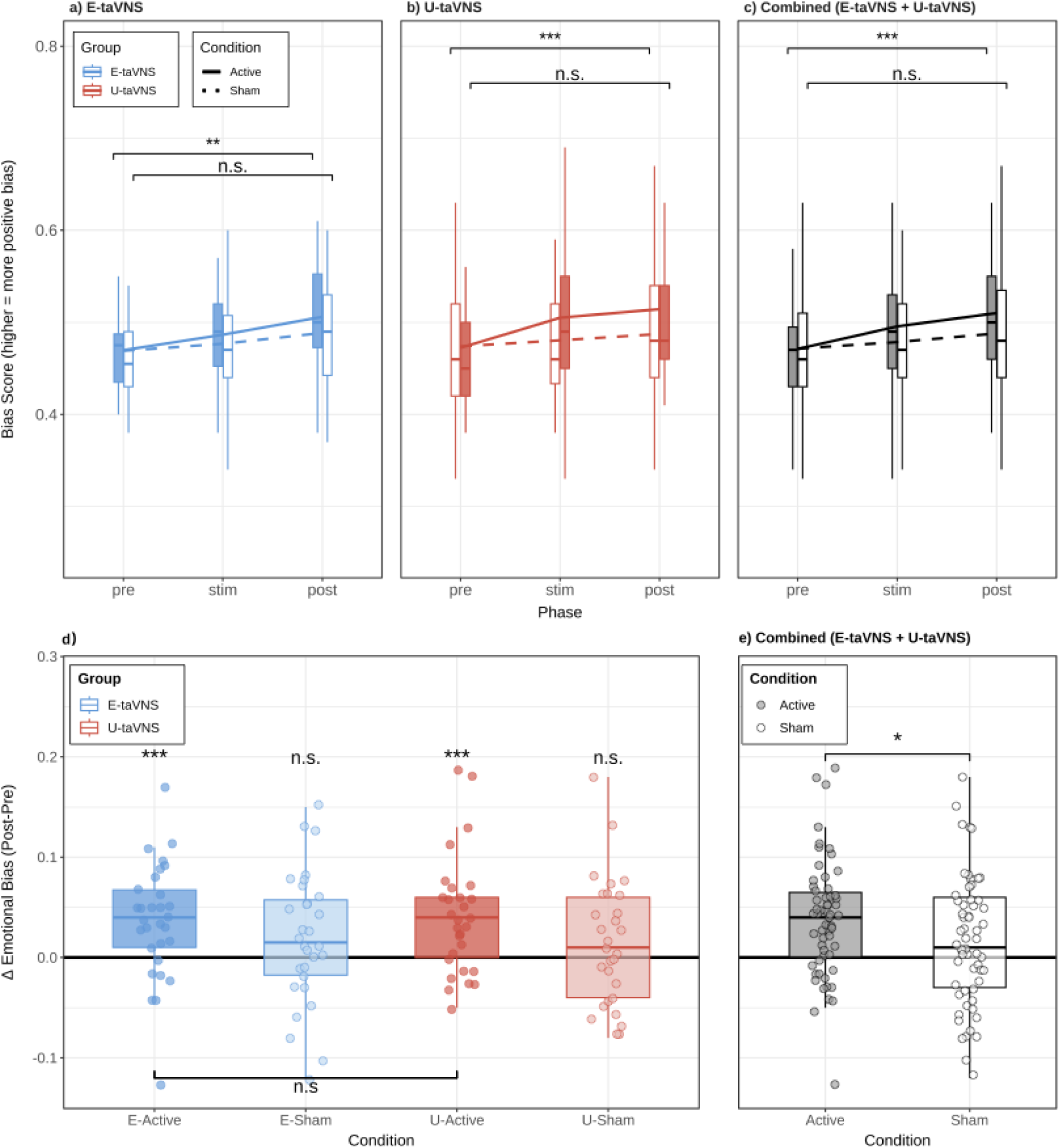
Effects of Electrical and Ultrasound taVNS on Emotional Bias. Emotional bias scores (proportion of faces categorized as “happy”) across (a) Pre, (b) During, and (c) Post phases for Active and Sham stimulation in the E-taVNS and U-taVNS groups. (d) Delta emotional bias scores (Post − Pre) for E-taVNS and U-taVNS separately under Active and Sham stimulation. (e) Combined analysis pooling both modalities, showing significantly greater positive bias shifts in the Active versus Sham condition. Dark colours are active and light colours are sham. * p < 0.05, ** p < 0.01, *** p < 0.001, n.s. not significant.

The individual-level data underlying these results are provided in Supplementary Figure 1.

To directly assess taVNS effects, delta bias scores (Post − Pre) were analyzed. Both active stimulation conditions significantly increased emotional bias scores (E-taVNS: t(29) = 3.51, p <.001, Cohen’s d = 0.640; U-taVNS: t(28) = 3.70, p <.001, Cohen’s d = 0.687) (Figure 3d). In contrast, sham stimulation did not produce significant changes (E-taVNS: t(29) = 1.59, p =.122, Cohen’s d = 0.290; U-taVNS: t(28) = 1.18, p =.248, Cohen’s d = 0.219). Within groups, paired-samples t-tests showed no significant differences between active and sham stimulation for E-taVNS, t(29) = −1.27, p =.107 (one-tailed), d = −0.23. The U-taVNS group showed a marginal effect, t(28) = −1.57, p =.063 (one-tailed), d = −0.29, with active stimulation trending toward stronger reductions in negative bias (Figure 3d).

To compare the efficacy between stimulation modalities, an independent samples t-test was conducted on the delta emotional bias scores between the E-taVNS and U-taVNS groups. Comparisons between modalities showed no significant differences in efficacy for either active (t(57) = −0.370, p =.712, Cohen’s d = −0.10) or sham (t(57) = 0.311, p =.757, Cohen’s d = 0.08) conditions (Figure 3d).

Given the absence of modality differences, both stimulation types were pooled to examine the overall effect of taVNS. The combined analysis revealed a significant main effect of Condition was observed, F(1,58) = 5.40, p =.024, 1]^2^ =.085, a significant main effect of Phase, F(2,116) = 14.59, p <.001, 1]^2^ =.201 (Figure 3c, Table S2). The Condition × Phase interaction showed the marginally significant effects, F(2,116) = 2.64, p =.076, 1]^2^ =.043. Post hoc comparisons confirmed a significant increase in bias score from the pre-to post-assessment in the active condition (mean difference = −0.039, p <.001), whereas no significant changes occurred in the sham condition (all ps >.34). Delta bias scores further supported this pattern, with active stimulation producing significantly greater reductions in negative bias than sham (t(58) = −2.04, p =.023 (one-tailed), d = −0.27) (Figure 3e).

### The relationship between taVNS effects and questionnaires

Linear regression analyses were conducted to examine whether interoceptive awareness, measured by the MAIA, predicted the emotional bias outcome. When using the MAIA total score as a predictor, the regression model was not statistically significant (B = 0.018, SE = 0.013, t = 1.38, p =.173). To explore potential differential contributions of specific interoceptive domains, separate regression models were run for each of the eight MAIA subscales. Only the Noticing subscale significantly predicted the effects of taVNS (B = 0.021, SE = 0.009, t = 2.45, p =.017), accounting for 9.6% of the variance in the outcome ( =.096) (Figure 4). The remaining seven subscales, Not-Distracting (p =.588), Not-Worrying (p =.436), Attention Regulation (p =.373), Emotional Awareness (p =.579), Self-Regulation (p =.851), Body Listening (p =.169), and Trusting (p =.376) did not significantly predict the outcome variable (Table S3). There were no significant correlations between BAI/BDI and taVNS effects.

**Figure 4.**
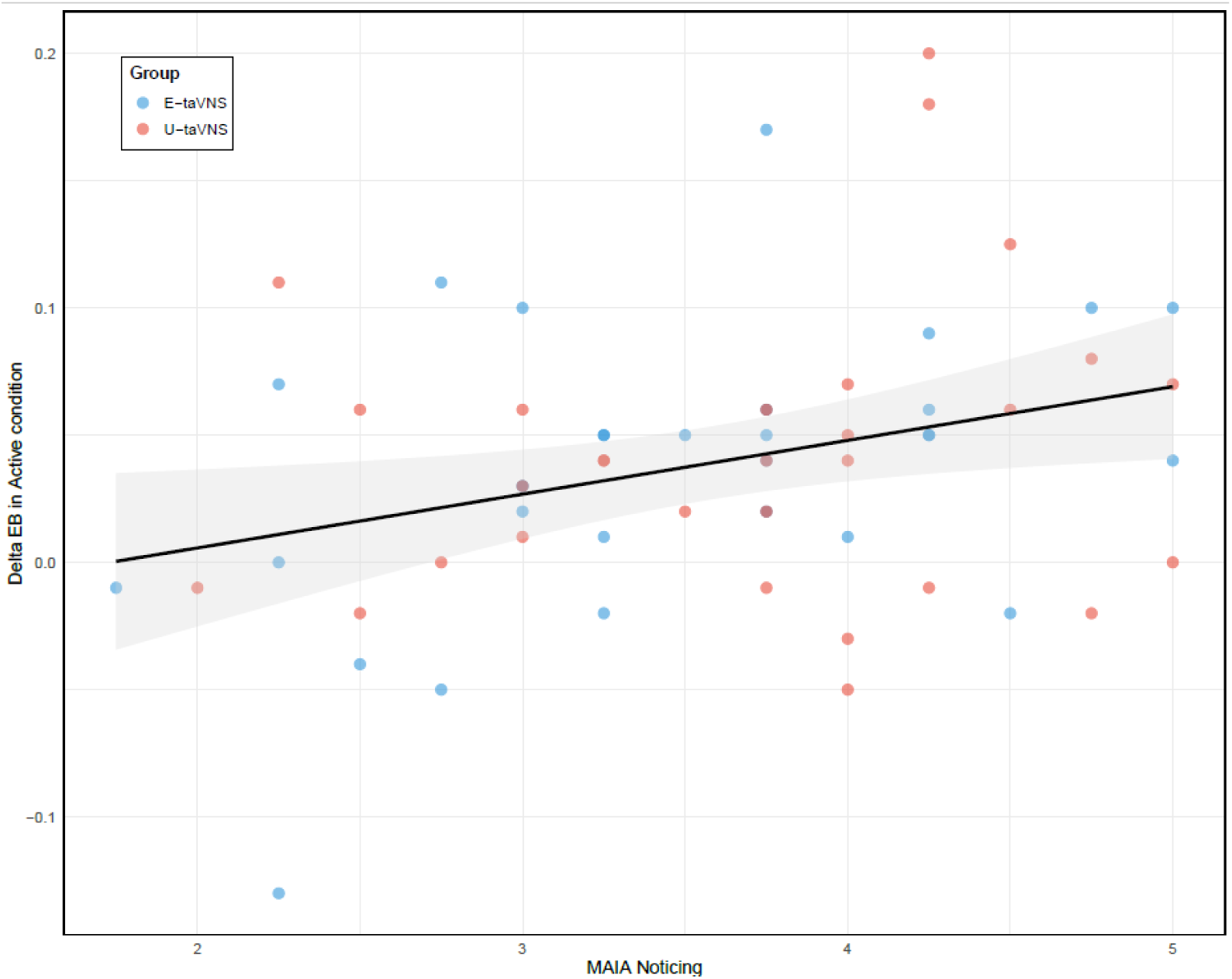
Relationship Between MAIA Noticing Scores and Change in Emotional Bias During Active Stimulation. Scatterplot illustrates the association between MAIA Noticing scores and delta emotional bias (Post − Pre) in the active stimulation condition. Each point represents an individual participant. Blue circles indicate participants in the E-taVNS group, and red circles indicate participants in the U-taVNS group. A higher Noticing score was associated with a greater shift toward positive emotional bias.

### Adverse Effects results

A chi-square test of independence was performed to examine the association between stimulation modality and the occurrence of aversive effects (Figure 5, Table S4). A higher proportion of participants in the E-VNS group reported skin irritation (20.0%) compared to those in the U-VNS group (3.4%) ( X^2^ = 3.86, p =.049), indicating U-taVNS was associated with a lower incidence of skin irritation than E-taVNS.

**Figure 5.**
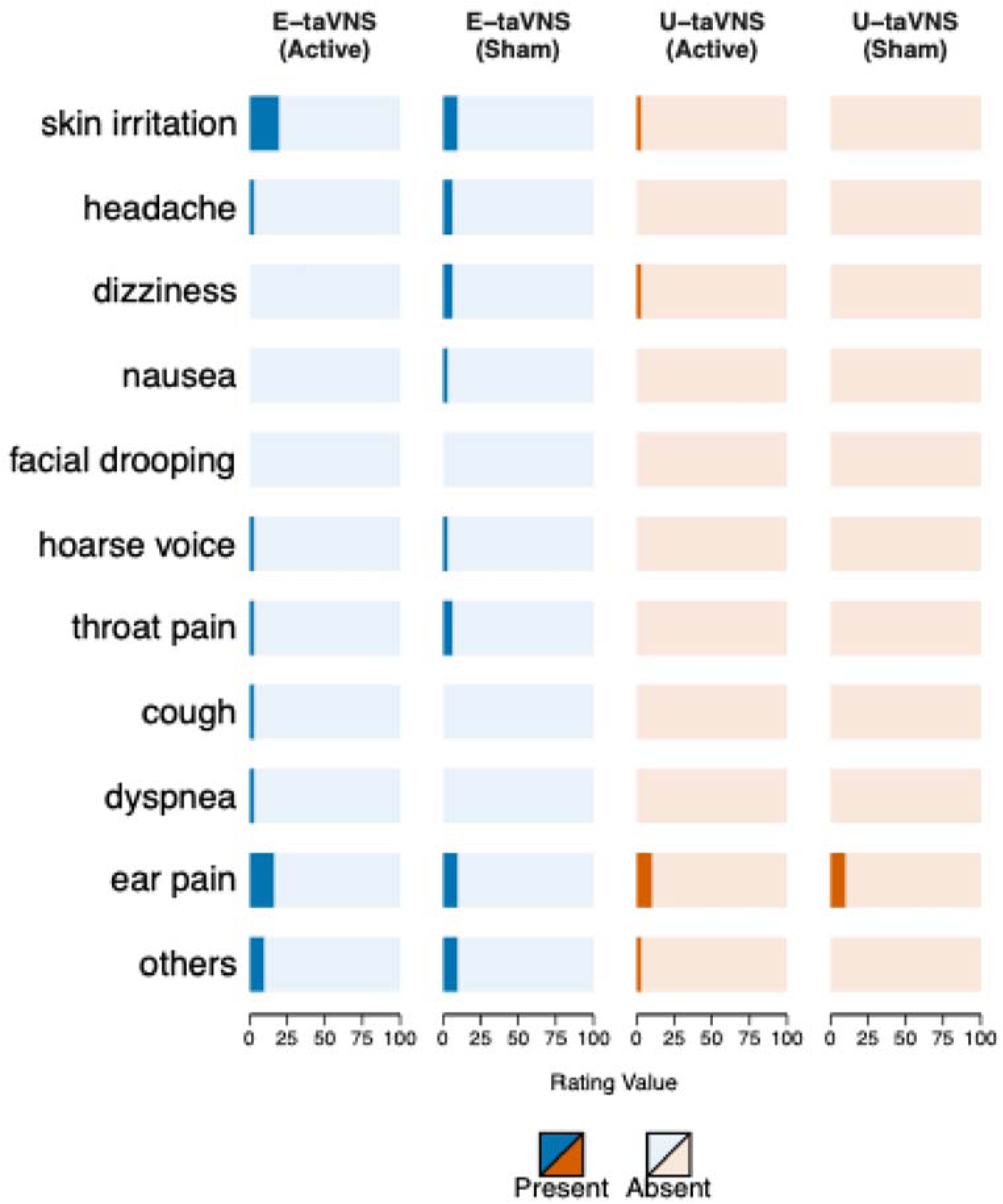
Reported Adverse Effects Across Electrical and Ultrasound taVNS Conditions. Visualization of the frequency of self-reported adverse effects during Active and Sham sessions for electrical taVNS (E-taVNS) and ultrasound taVNS (U-taVNS). Each bar represents the proportion of participants reporting the presence (colored bars) or absence (light bars) of each side effect, including skin irritation, headache, dizziness, nausea, facial drooping, hoarse voice, throat pain, cough, dyspnea, ear pain, and other symptoms. Blue shading indicates the E-taVNS group, and orange shading indicates the U-taVNS group.

## Discussion

The present study investigated the effects of both electrical and ultrasound transcutaneous vagus nerve stimulation (taVNS) on emotional bias in healthy young adults, using a facial emotion categorization task with morphed happy-sad faces. Our findings revealed that both stimulation modalities successfully shifted emotional bias in a positive direction, reflecting a reduction in negative bias. Consistent with our primary hypothesis, active taVNS significantly reduced negative emotional bias compared with both sham and baseline (pre-stimulation) conditions.

These findings align with previous research showing taVNS modulates affective processing and promotes positive mood states (Ferstl et al., 2022; Johnson & Steenbergen, 2022). The observed shift toward more positive interpretation of ambiguous facial expressions suggests that taVNS engages neural circuits underlying emotional categorization, potentially through enhanced LC-NE system activation and strengthened prefrontal-amygdala connectivity (Liu et al., 2016).

Moreover, the main effect of phase, along with post hoc evidence of bias reduction from pre-to post-stimulation, indicates that taVNS effects persist beyond the stimulation period.

The positive shift in emotional bias likely reflects the modulation of multiple interconnected systems. The vagus nerve’s afferent projections to the NTS and subsequent connections to the LC implicate noradrenergic mechanisms as primary mediators (Henry, 2002). Enhanced LC activity could improve cognitive flexibility and reduce attentional capture by negative stimuli, thereby facilitating more balanced emotional appraisal (Ghosh & Maunsell, 2024). Additionally, taVNS-induced changes in parasympathetic tone may contribute to emotional regulation by promoting physiological calmness and enhancing top-down control. Previous research has linked higher heart rate variability, a marker of parasympathetic activity, to reduced negativity bias in emotion perception (Osnes et al., 2023). The bidirectional communication between peripheral autonomic state and central emotional processing networks suggests that taVNS may induce a physiological state conducive to positive affective interpretation (Johnson & Steenbergen, 2022; Makovac et al., 2015). The comparable efficacy of both stimulation modalities further supports the idea that activation of ABVN afferents, rather than the specific method of activation, underlies the observed effects, highlighting the importance of anatomical targeting over stimulation modality. These findings are consistent with the well-established mechanisms of taVNS action. Afferent vagal projections from the auricular branch reach the NTS and subsequently the LC and amygdala–prefrontal circuits (Bonaz et al., 2018; Liu et al., 2016). By modulating the LC–norepinephrine system, taVNS enhances cognitive control and emotional regulation (Colzato & Beste, 2020; Ghosh & Maunsell, 2024). Increased parasympathetic tone and LC–prefrontal connectivity may jointly support adaptive reappraisal and reduced negativity bias (Koenig et al., 2021; Zhu et al., 2024). In addition to electrical stimulation, ultrasound-based taVNS may engage similar neural circuits through distinct biophysical mechanisms. Low-intensity focused ultrasound induces mechanical membrane deformation that influences neuronal excitability without direct electrical current (Dell’Italia et al., 2022; Yoo et al., 2022). Given its capacity to activate the same vagal afferent pathways projecting to the amygdala and hippocampus (Hacker et al., 2023; Kohler et al., 2025), U-taVNS may produce comparable affective modulation with more compliance and tolerability.

We found no significant differences in efficacy between electrical and ultrasound stimulation. Both E-taVNS and U-taVNS produced comparable shifts in emotional bias, suggesting that these distinct physical modalities converge on shared neurophysiological pathways. This equivalence is particularly noteworthy given the fundamental differences in mechanisms, electrical stimulation involves direct current-induced depolarization, whereas ultrasound stimulation relies on mechanical pressure waves that deform neuronal membranes and influence excitability (Dell’Italia et al., 2022; Yoo et al., 2022). An important consideration, however, is that ultrasound in this study was applied only unilaterally. Thus, while both modalities showed similar outcomes, this comparison should be interpreted with caution, as ultrasound stimulation might have yielded stronger effects had it been delivered bilaterally. This point is particularly relevant given that our electrical taVNS device stimulated both sides, whereas most conventional taVNS approaches target only the left ear due to ongoing debate about potential right-sided cardiac influences. Although efficacy was similar, U-taVNS demonstrated greater tolerability.

Only 3.4% of participants in the ultrasound group reported skin irritation compared with 20.0% in the electrical group, suggesting that ultrasound stimulation may be a more tolerable alternative for long-term therapeutic applications. This improved tolerability could enhance treatment adherence, particularly important for conditions requiring extended stimulation protocols such as depression or anxiety disorders (Bretherton et al., 2019; Tan et al., 2023)

While the MAIA total score did not significantly predict the changes in emotional bias, the Noticing subscale, reflecting awareness of bodily sensations, emerged as a significant predictor. This finding highlights the potential relevance of interoceptive awareness in mediating taVNS effects on emotion processing, consistent with evidence linking heightened body awareness to adaptive emotional regulation (Hanley et al., 2017, Zamariola et al., 2019).

Several limitations should be considered when interpreting our results. First, our sample consisted of healthy young adults, limiting generalizability to clinical populations where emotional biases are more pronounced. Second, we examined only acute effects; whether these changes persist or accumulate with repeated stimulation remains unknown. Third, the absence of physiological measures (e.g., heart rate variability, pupillometry, or salivary biomarkers) limits our ability to confirm proposed mechanistic pathways. Finally, our emotional bias task utilized a limited set of facial identities and emotions (happy and sad); future studies should employ more diverse stimuli to ensure generalizability across different emotional categories and intensities.

Despite these limitations, the findings carry meaningful clinical implications. Demonstrating that both taVNS modalities can shift emotional bias in healthy individuals provides proof-of-principle for potential therapeutic applications in mood and anxiety disorders, where negative bias is a core maintaining factor (Mathews & MacLeod, 2005). The superior tolerability of U-taVNS, combined with comparable efficacy, positions it as a promising candidate for home-based and long-term therapeutic protocols.

In conclusion, this study provides evidence that both electrical and ultrasound-based taVNS can acutely modulate emotional bias in healthy adults. These findings advance our understanding of non-invasive vagus nerve stimulation as a tool for modulating affective processing. As the field moves toward clinical implementation, our results support continued investigation of taVNS approaches for treating conditions characterized by maladaptive emotional biases.

## Acknowledgments

The authors acknowledge all participants.

## Author contributions

Conceptualization: JJ, KJ; Methodology: JJ, MK, HC, JS; Investigation: JJ, JYN; Writing— JJ, KJ, JYN; Writing—review & editing: JJ, KJ, JYN, MK, HC, JS

