## Supplementary Information for "Transcutaneous auricular vagus nerve stimulation enhances emotional bias towards happiness in healthy young adults: A comparative study of electrical and ultrasound stimulation"

Supplementary Figure 1

Supplementary Table 1

Supplementary Table 2

Supplementary Table 3

Supplementary Table 4

**Supplementary Figure 1**

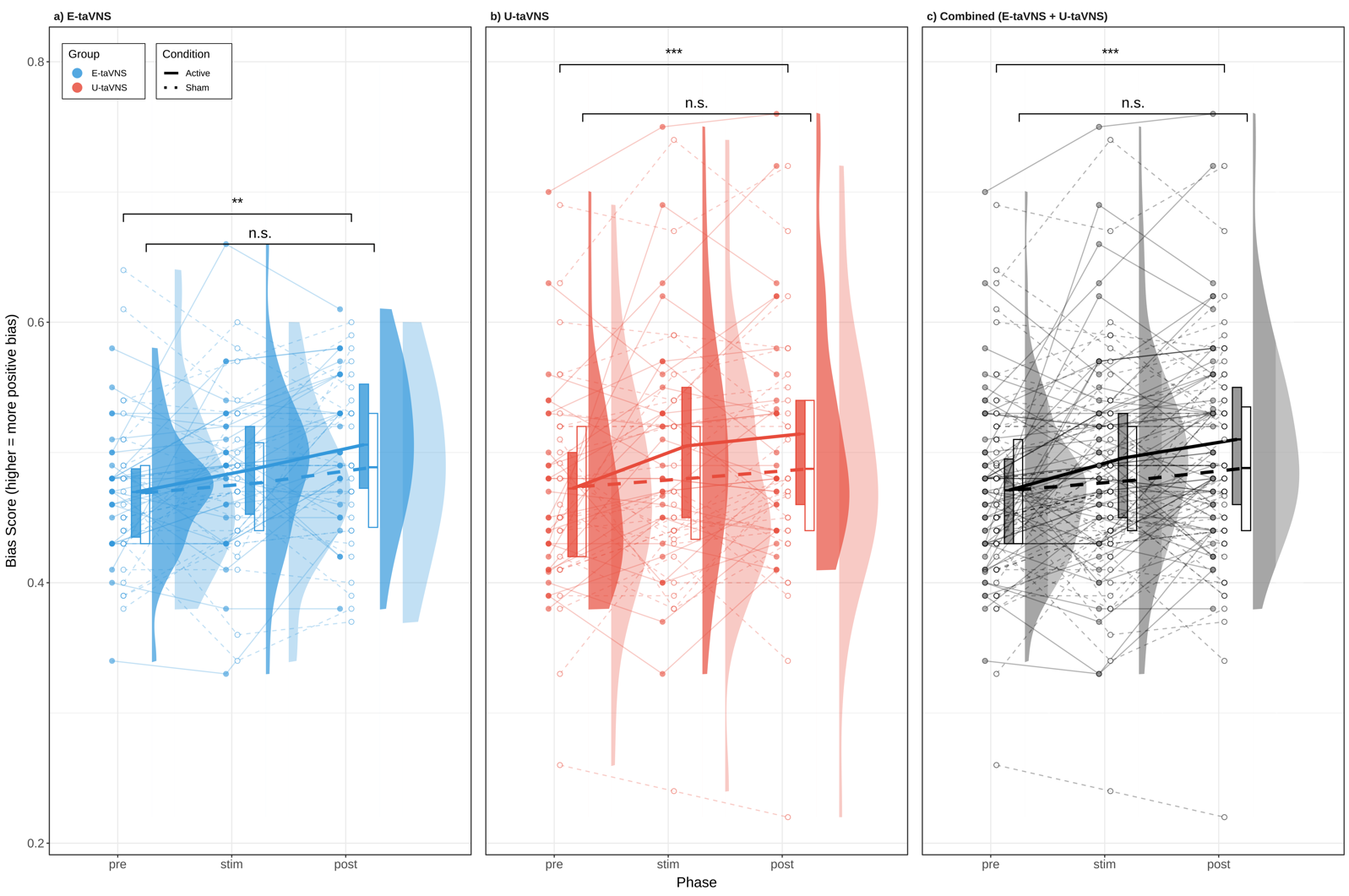

**Figure S1.** Effects of Electrical and Ultrasound taVNS on Emotional Bias, raincloud plot with individual data points. *** p < 0.001**,** ** p < 0.01, * p < 0.05, n.s. not significant.

**Supplementary Table 1**

| Effect | $df_{1}$ | $df_{2}$ | $F$ | $p$ | $\eta_{p}^{2}$ |
| --- | --- | --- | --- | --- | --- |
| Group | 1 | 57 | 0.13 | .716 | .002 |
| Condition | 1 | 57 | 5.40 | .024*^*^* | .008 |
| Phase | 2 | 114 | 14.43 | $<$.001*^***^* | .023 |
| Group $\times$ Condition | 1 | 57 | 0.43 | .516 | .001 |
| Group $\times$ Phase | 2 | 114 | 0.34 | .712 | .001 |
| Condition $\times$ Phase | 2 | 114 | 2.64 | .076 | .004 |
| Group $\times$ Condition $\times$ Phase | 2 | 114 | 0.35 | .708 | .001 |

Table S1. Mixed-design ANOVA results. *** p < 0.001**,** ** p < 0.01, * p < 0.05.

**Supplementary Table 2**

| Effect | $df_{1}$ | $df_{2}$ | $F$ | $p$ | $\eta_{p}^{2}$ |
| --- | --- | --- | --- | --- | --- |
| Condition | 1 | 58 | 5.40 | .024 | .085 |
| Phase | 2 | 116 | 14.59 | $<$.001 | .201 |
| Condition $\times$ Phase | 2 | 116 | 2.64 | .076 | .043 |

Table S2. Mixed-design ANOVA results (collapsed across Group) $\eta_{p}^{2}$ = partial eta squared. . *** p < 0.001**,** ** p < 0.01, * p < 0.05.

**Supplementary Table 3**

| **Predictor** | **B** | **SE** | **t** | **p** | $R^{2}$ |
| --- | --- | --- | --- | --- | --- |
| MAIA Total Score | 0.018 | 0.013 | 1.38 | .173 | .032 |
| Noticing | 0.021 | 0.009 | 2.45 | .017* | .096 |
| Not-Distracting | 0.004 | 0.008 | 0.55 | .588 | .005 |
| Not-Worrying | $-$0.006 | 0.007 | $-$0.78 | .436 | .011 |
| Attention Regulation | 0.007 | 0.008 | 0.90 | .373 | .014 |
| Emotional Awareness | 0.005 | 0.009 | 0.56 | .579 | .005 |
| Self-Regulation | 0.001 | 0.007 | 0.19 | .851 | .001 |
| Body Listening | 0.007 | 0.005 | 1.39 | .169 | .033 |
| Trusting | 0.006 | 0.006 | 0.89 | .376 | .014 |

Table S3 Relationship Between MAIA Scores and Combined taVNS Efficacy (Linear Regression Model), MAIA = Multidimensional Assessment of Interoceptive Awareness, $\eta_{p}^{2}$ = partial eta squared. *** p < 0.001**,** ** p < 0.01, * p < 0.05.

**Supplementary Table 4**

| **Adverse Effect** | **E-taVNS** | **U-taVNS** | $\chi^{2}$ | **p-value** |
| --- | --- | --- | --- | --- |
|  | (n=30) | (n=29) |  |  |
| Skin Irritation | 6 (20%) | 1 (3.4%) | 3.86 | 0.049* |
| Headache | 1 (3.3%) | 0 (0%) | 0.98 | 0.321 |
| Dizziness | 0 (0%) | 1 (3.4%) | 1.05 | 0.305 |
| Nausea/Vomiting | 0 (0%) | 0 (0%) | - | - |
| Facial Drooping | 0 (0%) | 0 (0%) | - | - |
| Hoarse Voice | 1 (3.3%) | 0 (0%) | 0.98 | 0.321 |
| Throat/Neck Pain | 1 (3.3%) | 0 (0%) | 0.98 | 0.321 |
| Cough | 1 (3.3%) | 0 (0%) | 0.98 | 0.321 |
| Shortness of Breath | 1 (3.3%) | 0 (0%) | 0.98 | 0.321 |
| Ear Pain | 5 (16.7%) | 3 (10.3%) | 0.50 | 0.478 |
| Others | 3 (10%) | 1 (3.4%) | 1.00 | 0.317 |

Table S4 Adverse Effects During Active Stimulation by Group. Values shown as n (%). Chi-square test with df=1. *** p < 0.001**,** ** p < 0.01, * p < 0.05.
